# A role for phagocytosis in inducing cell death during thymocyte negative selection

**DOI:** 10.1101/624148

**Authors:** Nadia S. Kurd, Lydia K. Lutes, Jaewon Yoon, Ivan L. Dzhagalov, Ashley Hoover, Ellen A. Robey

## Abstract

Autoreactive thymocytes are eliminated during negative selection in the thymus, a process important for establishing self-tolerance. Thymic phagocytes serve to remove dead thymocytes, but whether they play additional roles during negative selection remains unclear. Here, we demonstrate that phagocytosis promotes negative selection, and that negative selection is more efficient when the phagocyte also presents the negative selecting peptide. Our findings support a two-step model for negative selection in which thymocytes initiate the death process following strong TCR signaling, but ultimately depend upon phagocytosis for their timely death. Thus, the phagocytic capability of cells that present self-peptides is a key determinant of thymocyte fate.

## Introduction

During negative selection, thymocytes bearing self-reactive T cell receptors (TCR) are eliminated from the T cell repertoire, an important process for the establishment of self-tolerance. Thymocytes interact with a variety of thymic-resident cells that present self-peptide:MHC complexes, and thymocytes bearing TCRs with high affinity for self-ligands can receive apoptotic death signals^1^. Apoptosis is an immunologically silent form of cell death that is generally thought to be cell-autonomous once initiated^2^. Although peptide-presenting cells provide the initial apoptotic stimulus to autoreactive thymocytes, whether additional cellular interactions are required to mediate thymocyte death remains unknown.

In addition to the affinity of TCR for self-peptide-MHC, the nature of the peptide-presenting cell is also an important determinant of T cell fate. For example, hematopoietic cells, especially dendritic cells (DC), are potent inducers of negative selection, whereas cortical thymic epithelial cells (cTEC) are specialized to mediate positive selection, promoting thymocyte maturation and survival ^3–8^. Distinctive features of these cell types, including specialized peptide processing machinery in cTECs and high expression of costimulatory ligands in DCs, play an important role in instructing divergent thymocyte fates^1,8^. Peptide repertoire and costimulation can contribute to the intensity of TCR signal experienced by a thymocyte, but whether the peptide-presenting cells provide additional TCR-independent signals to promote thymocyte death or differentiation is largely unknown.

A related question is whether a strong TCR signal is sufficient to commit thymocytes to die in a cell-autonomous fashion, or whether other cellular interactions are required. Early observations of apoptotic bodies within thymic phagocytes ^2,9^, together with time-lapse microscopy of thymocytes undergoing negative selection^3–8,10^, suggest a close coupling between thymocyte death and phagocytosis. In particular, visualization of dying thymocytes and phagocytes within thymic tissue slices showed that the death of autoreactive thymocytes invariably occurred during close contact with phagocytes, and in most cases death appeared to occur after phagocytosis^10^. However, it remained unclear whether phagocytes actually contributed to the death of thymocytes by serving as peptide-presenting cells and/or by actively inducing cell death.

Dendritic cells are the most potent antigen-presenting cells (APC) for priming naïve T cells and also present self-peptide for negative selection^3–7,11^, whereas thymic macrophages are known for their role in clearing away apoptotic thymocytes^9^. Nevertheless, there is considerable functional and phenotypic overlap between DC and macrophages. For example, the marker F4/80 is often used to identify macrophages, but also marks a population with substantial antigen presentation function^12^. Likewise, the marker CD11c is often used to identify DCs, but is co-expressed with F4/80 by a subset of DC-like cells in the thymus^13^. To what extent the functions of peptide presentation and phagocytic clearance reside within separate or overlapping thymic cell populations remains unclear.

Phagocytes recognize and uptake apoptotic cells via receptors for “eat-me” signals displayed on the surface of dying cells ^14^. For example, apoptosis induces asymmetry in the plasma membrane, leading to the exposure of phosphatidylserine (PS), which is then recognized by PS receptors on phagocytes. A variety of “eat-me” receptors are expressed and functional in the thymus^15–17^, but the mechanisms that mediate the efficient removal of autoreactive thymocytes during negative selection have not yet been clearly defined.

Here, we used a thymic slice system in which thymocytes undergo negative selection *in situ* to address these questions. We found that depletion of thymic phagocytes or blocking phagocytosis impaired negative selection, allowing for the increased survival of thymocytes that had experienced strong TCR signals. We also identified the PS receptor Tim-4 as an important player during negative selection to tissue-restricted antigens (TRA). Finally, we demonstrated that negative selection is most efficient when the same cell both presents the agonist peptide, and phagocytoses the self-reactive thymocyte. Taken together, our data suggest a two-step model for negative selection in which strong TCR signals initiate the apoptotic program, but thymocytes depend on phagocytosis for their timely death. Thus, thymic phagocytes are not merely “scavengers”, but rather play prominent roles in the induction of self-tolerance, both as peptide-presenting cells and as active inducers of self-reactive thymocyte death.

## Results

### Depletion of thymic phagocytes inhibits negative selection

To examine the role of phagocytes during negative selection, we used thymic tissue slices prepared from Macrophage-Associated Fas-Induced Apoptosis (MAFIA) mice. In these mice, an inducible suicide gene under the control of the colony stimulating factor 1 receptor (Csf1R) promoter is expressed in DC and macrophage subsets^18^. We have previously observed closely coupled thymocyte death and phagocytosis by GFP^+^ cells in thymic slices from LysMGFP reporter mice ^10,19^. Flow cytometric analysis of the thymus of LysMGFP mice revealed that GFP-expressing cells include a subset of F4/80^hi^ macrophages that have been previously described as having potent phagocytic abilities, as well as a subset of CD11c^hi^ DCs (Supplementary Fig. 1a)^10,16^. We confirmed expression of the MAFIA transgene in these subsets using flow cytometric analysis to measure expression of a linked GFP gene (Supplementary Fig. 2a). We observed that treatment of thymic slices for 16 hours of culture with the small molecule inducer AP20187 led to the depletion of approximately 50% of F4/80^hi^ macrophages and CD11c^hi^ DC (Supplementary Fig. 2b).

To assess the impact of phagocyte depletion on negative selection, we used a previously described peptide-induced model of negative selection^10,20^. Thymocytes from mice with a defined MHC class I-restricted TCR transgene (OT-I) were overlaid onto thymic slices with or without the cognate antigen (SIINFEKL, OVAp), along with a reference thymocyte population (F5 TCR transgenic or polyclonal WT thymocytes) (Fig. 1a). We used flow cytometry to determine the ratio of viable OT-I:reference thymocytes remaining in the slice as a measure of negative selection ^10,20,21^ (Fig. 1a). Consistent with our previous study^10^, we observed a substantial reduction in the number of live OT-I thymocytes relative to reference thymocytes in thymic slices containing OVAp after 16 hours of culture (Fig. 1b). In contrast, on phagocyte-depleted slices containing OVAp, the ratio of live OT-I:reference thymocytes was similar to that in -OVAp control slices, consistent with the idea that phagocytes promote negative selection.

**Figure 1.**
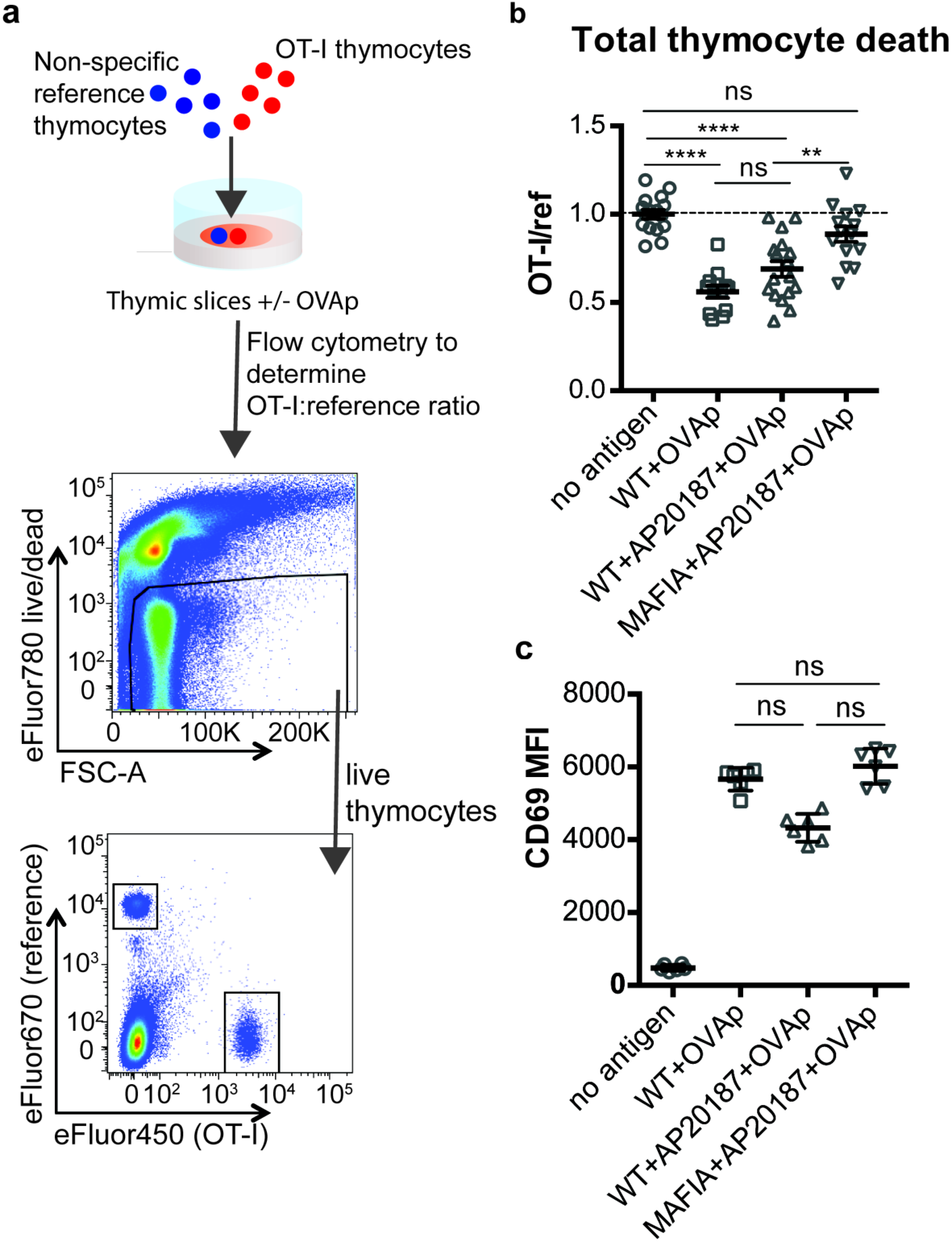
Depletion of phagocytes inhibits negative selection without dampening antigen recognition. (a) Strategy to quantify negative selection: labeled OT-I and reference thymocytes (either wild type or F5 TCR transgenic) were overlaid onto thymic slices with or without OVA peptide, cultured for 16 hours, and then dissociated for analysis by flow cytometry. Lower panels show the flow cytometry gating strategy used to quantify live OT-I and reference thymocytes. (b-c) For depletion of phagocytes, WT or MAFIA thymic slices were treated with AP20187 and cultured for an additional 16 hours prior to the addition of thymocytes and peptide. b) Thymocyte death displayed as the ratio of live OT-I thymocytes relative to live reference thymocytes present within the slice. We further normalized the ratios of OT-I:reference thymocytes in each experiment so that the average of the corresponding “no OVA” samples was 1.0. (c) Expression of the activation marker CD69 by OT-I thymocytes displayed as Mean Fluorescence Intensity (MFI). ns not significant (p>0.05), **p<0.01, ****p<0.0001 (one-way ANOVA with Bonferroni’s correction with a 95% confidence interval, b, or Kruskal-Wallis test with Dunn’s multiple comparisons with a 95% confidence interval, c). Data are pooled from 3 independent experiments (b), or representative of 3 independent experiments (c), with mean and SEM of n=16 (b) or 5 (c) total slices per condition, where each dot represents an individual slice.

Thymic phagocytes, including DCs, serve as APCs during negative selection^4,6,8,12^. This raises the possibility that the observed defect in negative selection on phagocyte-depleted slices could be the result of thymocytes receiving insufficient TCR signals. However, OT-I thymocytes on phagocyte-depleted slices did not show decreased levels of the TCR activation marker CD69, arguing against impairment of TCR signals (Fig. 1c). These results suggest that thymocytes on phagocyte-depleted thymic slices exhibit enhanced survival despite continuing to receive strong TCR signals.

### Negative selection and phagocytosis in a model of tissue-restricted antigen presentation

Addition of agonist peptide directly to thymic tissue slices serves as a model for negative selection to ubiquitous self-antigen. To further characterize the relationship between thymocyte death and phagocytosis, we also examined a model of negative selection to TRA. In RIPmOVA transgenic mice, the model antigen ovalbumin (OVA), which provides the agonist peptide for OT-I thymocytes, is expressed in a subset of medullary thymic epithelial cells (mTECs), and is presented in the medulla by mTECs and hematopoietic-derived cells^3,22^. Because approximately 50% of CD4^+^CD8^+^ double positive thymocytes, and all of the more mature CD8^+^ single positive thymocytes from OT-I mice express a medullary chemokine receptor pattern (CXCR4^−^CCR7^+^) and migrate to the medulla in thymic slices^23–26^, we expected that the majority of OT-I thymocytes would encounter OVA when added to RIPmOVA slices. We observed that approximately 50% of OT-I thymocytes were lost by 9 hours, and there was no further reduction through 48 hours of culture (Fig. 2a). Thus, the timing and extent of negative selection in a TRA model are in line with our previous results using a ubiquitous model of negative selection.

**Figure 2.**
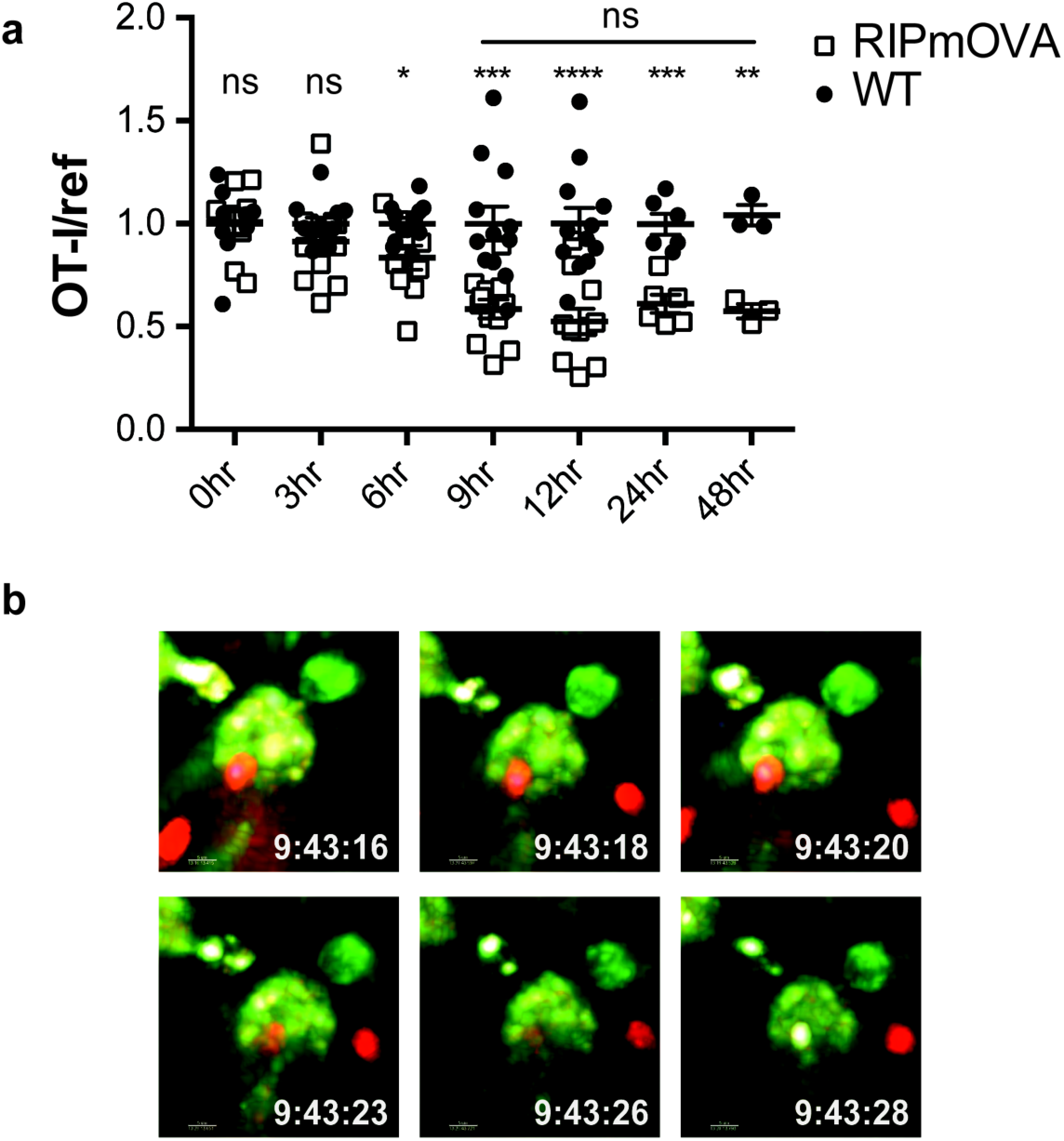
Thymocyte death and phagocytosis during negative selection to tissue-restricted antigen. Negative selection on RIPmOVA slices (open squares), displayed as the ratio of live OT-I thymocytes relative to live reference thymocytes present within the slice, normalized to no antigen controls (black squares). Data are pooled from 4 (0, 3, 6, 9, and 12 hour timepoints), 2 (24 hour timepoint), or 1 (48 hour timepoint) experiments, with mean and SEM of n=3 (48hr WT and RIPmOVA), 6 (24hr WT and RIPmOVA), 9 (0hr WT and RIPmOVA), 7 (6hr RIPmOVA), 11 (12hr WT), 12 (3hr and 9hr WT and RIPmOVA, 6hr and 12hr WT) total slices per condition, respectively, where each dot represents an individual slice. ns not significant (p>0.05), *p<0.05, **p<0.01, ***p<0.001, ****p<0.0001. Unpaired two-tailed Student’s *t*-test of WT vs RIPmOVA for each timepoint, or two-way ANOVA with 95% confidence interval with Tukey’s multiple comparisons test to compare RIPmOVA samples across timepoints (horizontal line). (b) Still images from a time-lapse series showing an example of OT-I thymocyte death. OT-I thymocytes were depleted of mature CD8 SP and labeled with Hoechst and SNARF before overlaying on LysMGFP RIPmOVA thymic slices. Slices were imaged by two-photon scanning laser microscopy. A 30-minute movie was recorded in the medulla, with the time elapsed since thymocyte entry into the slice shown in white.

We used 2-photon time-lapse microscopy to visualize the interactions between OT-I thymocytes and phagocytes on RIPmOVA thymic slices^10^. To visualize thymocyte death, we used a previously described method in which thymocytes are double-labeled with a cytosolic dye (SNARF, shown in red) that escapes as cells lose membrane integrity, and a nuclear dye (Hoechst, shown in blue) that increases signal during apoptosis-associated chromatin changes^10,27^. OT-I thymocytes were overlaid onto thymic slices from LysMGFP RIPmOVA transgenic mice and imaged after 6-10 hours of culture. Consistent with our earlier study^10^, cell death occurred while thymocytes were in close contact with, or engulfed by, GFP-expressing phagocytes (Fig. 2b, Supplementary Movies 1-2). Thus, the close coupling of autoreactive thymocyte death and phagocytosis occurs during negative selection to both tissue-restricted and ubiquitous self-antigens.

### Phagocyte killing of autoreactive thymocytes is mediated by phosphatidyl serine receptors

To determine whether phagocytes exert their effect on negative selection specifically through phagocytosis, we evaluated the efficiency of negative selection in an environment in which phagocytes are present, but impaired in their ability to phagocytose. Phosphatidylserine (PS) is exposed at the cell surface early in the apoptotic process, and serves as an “eat-me” signal to phagocytes^14^. To determine whether PS mediates phagocytosis of self-reactive thymocytes, we used Annexin V (AnnV), a small protein that binds to PS, to block the interaction between PS and its receptors in thymic slices^28^. Negative selection on RIPmOVA thymic slices was completely blocked by treatment of thymic slices with AnnV (Fig. 3a), while antigen recognition, as assessed by CD69 upregulation, was not impaired (Fig. 3b). AnnV addition had a similar effect on negative selection in response to OVAp (Fig. 3c,d). These results confirm that phagocytosis promotes the death of autoreactive thymocytes, and suggest that PS receptors are important for this process.

**Figure 3.**
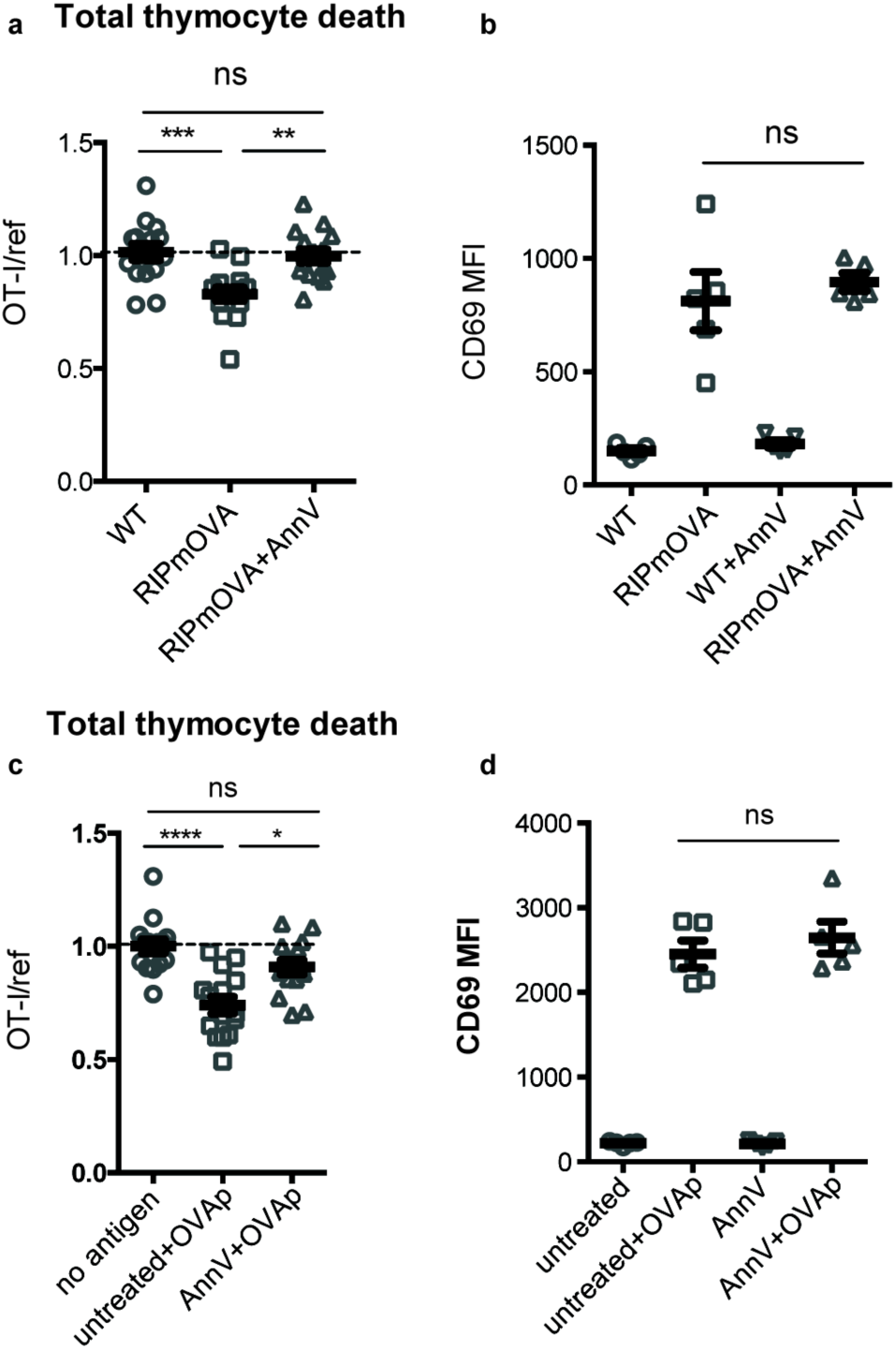
Phagocyte killing of autoreactive thymocytes is mediated by phosphatidylserine receptors. OT-I and reference thymocytes in AnnV buffer with or without AnnV were overlaid onto RIPmOVA slices (a,b) or WT slices treated with OVAp (c,d). Slices were then treated with AnnV and harvested 16 hours later for flow cytometric analysis. (a,c) Negative selection displayed as the ratio of live OT-I thymocytes relative to live reference thymocytes, normalized to no antigen controls. (b,d) Antigen recognition of OT-I thymocytes displayed as Mean Fluorescence Intensity (MFI) of the activation marker CD69. ns not significant (p>0.05), **p<0.01, ***p<0.001, ****p<0.0001 (one-way ANOVA with Bonferroni’s correction with a 95% confidence interval, a,c,d, or two-tailed unpaired Mann-Whitney test with 95% confidence interval, b). Data are pooled from 3 independent experiments (a,c), or representative of 3 independent experiments (b,d), with mean and SEM of n=15 (a,c) or 5 (b,d) total slices per condition, where each dot represents an individual slice.

### The phosphatidylserine receptor Tim-4 promotes negative selection of CD8 T cells in the thymus

Phagocytes express a number of receptors for PS, allowing them to recognize and uptake apoptotic cells^14^. These include Tim-4, a PS receptor previously reported to be expressed and functional in the thymus^15^. Using flow cytometry, we found that Tim-4 is expressed by almost all F4/80^hi^ macrophages and ∼30% of CD11c^hi^ DCs (Supplementary Fig. 3a). To investigate whether Tim-4 plays a role in negative selection, we prepared thymic tissue slices from Tim-4^−/−^ RIPmOVA transgenic mice, overlaid OT-I and reference thymocytes, and assessed the extent of negative selection after 16 hours of culture. OT-I thymocyte death was not detectable in Tim-4^−/−^ RIPmOVA thymic slices, but was readily detectable in thymic slices from age and sex-matched Tim-4^+/+^ RIPmOVA controls (Fig. 4a). Negative selection in response to OVAp was also slightly reduced on Tim-4^−/−^ thymic slices, although the difference did not reach statistical significance (Fig. 4c). Normal CD69 upregulation by OT-I thymocytes suggested that OVA is presented normally in the Tim-4^−/−^ thymic environment (Fig. 4b,d). This is consistent with the normal number and phenotype of thymic phagocytes that we observed in Tim-4^−/−^ mice (Supplementary Fig. 3b,c,d). Taken together, these data support the idea that phagocytosis is required for effective negative selection, and suggest that Tim-4 is a relevant player in this process.

**Figure 4.**
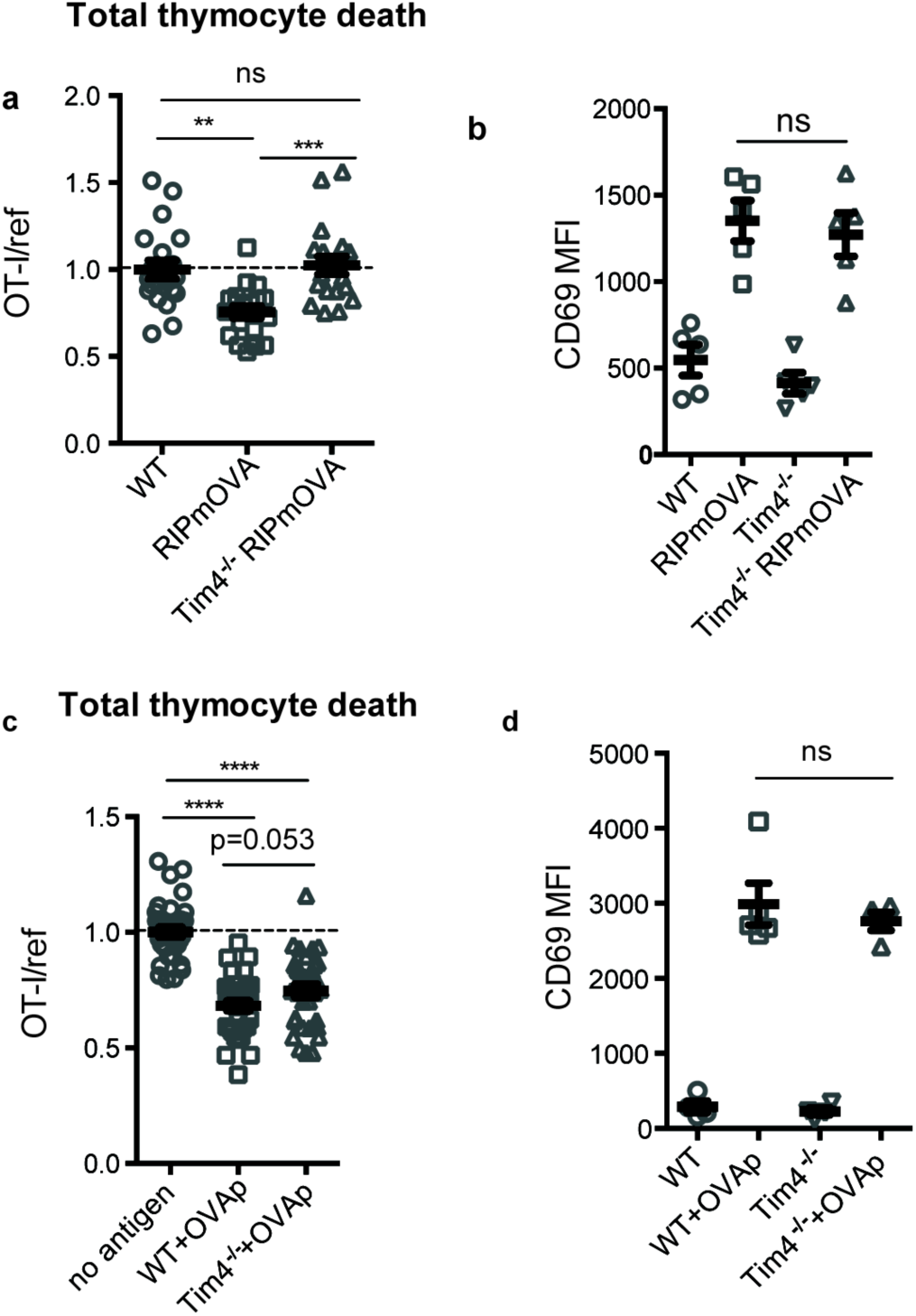
The phosphatidylserine receptor Tim-4 promotes negative selection to tissue-restricted antigens. OT-I and reference thymocytes were overlaid onto WT or Tim-4^−/−^ thymic slices with or without the RIPmOVA transgene (a,b), or with or without addition of OVAp (c,d), and slices were dissociated and analyzed by flow cytometry 16 hours later. (a,c) Negative selection displayed as the ratio of surviving OT-I thymocytes relative to reference thymocytes, normalized to no antigen controls. (b,d) Antigen recognition of OT-I thymocytes displayed as Mean Fluorescence Intensity (MFI) of the activation marker CD69. ns not significant (p>0.05), **p<0.01, ***p<0.001, ****p<0.0001 (one-way ANOVA with Bonferroni’s correction with 95% confidence interval, a-c, or Kruskal-Wallis test with Dunn’s multiple comparisons with 95% confidence interval, d). Data are pooled from 4 (a) or 7 (c) independent experiments, or representative of 4 (b) or 7 (d) independent experiments, with mean and SEM of n=20 (a), 35 (c) or 5 (b,d) total slices per condition, where each dot represents an individual slice.

### Antigen presentation by phagocytes promotes efficient negative selection

The close association between autoreactive thymocytes and phagocytes just prior to their engulfment and death^10^ (Fig. 2c) suggests that phagocytes may also serve as APCs. Moreover, the ability of a peptide-presenting cell to phagocytose may make it more potent at inducing negative selection. To test this idea, we took advantage of the fact that bone marrow-derived dendritic cells (DCs) have phagocytic activity (Supplementary Fig. 4) and can serve as exogenous peptide presenting cells when overlaid and allowed to migrate into thymic slices ^29,30^. We added OVA-loaded Tim-4^−/−^ or WT DCs onto thymic slices that had been previously overlaid with OT-I and reference thymocytes, and measured the extent of negative selection 16 hours later (Fig. 5a). Interestingly, while OVA-loaded WT DCs induced a significant level of negative selection, Tim-4^−/−^ DCs failed to induce detectable negative selection (Fig. 5b). This defect was not due to a deficiency in peptide presentation by Tim-4^−/−^ DCs, as there was no significant difference in the upregulation of CD69 by OT-I thymocytes (Fig. 5c). The fact that peptide-presenting cells defective in phagocytosis were unable to effectively induce negative selection, despite the fact that functional (non-presenting) endogenous phagocytes were present in the thymic slice, indicates that negative selection is more efficient when the same cell serves both as APC and phagocyte.

**Figure 5.**
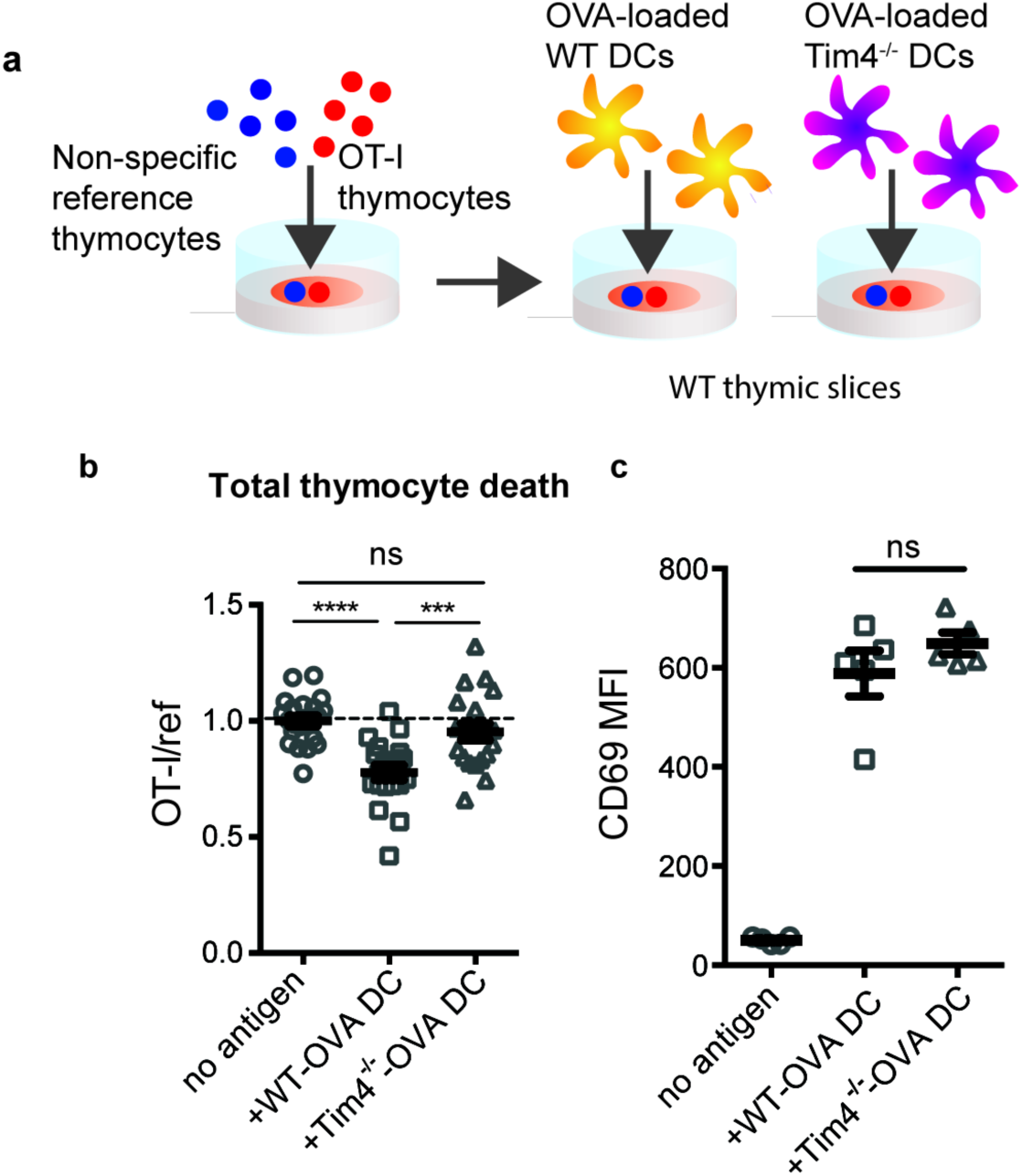
Peptide presentation by phagocytes promotes efficient negative selection. (a) Schematic of the experimental setup: OT-I and reference thymocytes were overlaid onto WT thymic slices onto which OVA-loaded or unloaded WT or Tim-4^−/−^ BMDCs were added. Slices were dissociated and analyzed by flow cytometry 16 hours later. (b) Negative selection displayed as the ratio of live OT-I thymocytes relative to live reference thymocytes, normalized to no antigen controls. Data are pooled from 4 experiments, with mean and SEM of n=20 total slices per condition shown in red. (c) Antigen recognition of OT-I thymocytes displayed as Mean Fluorescence Intensity (MFI) of the activation marker CD69. Data are representative of 4 independent experiments. ns not significant (p>0.05), ***p<0.001, ****p<0.0001 (one-way ANOVA with Bonferroni’s correction with 95% confidence interval). Data are pooled from 4 independent experiments (b), or representative of 4 independent experiments (c), with mean and SEM of n=20 (a), or 5 (c) total slices per condition, where each dot represents an individual slice.

## Discussion

While the importance of phagocytes in clearing dead thymocytes has long been appreciated, their role during negative selection prior to thymocyte death remained unknown. Here, we provide evidence for key roles for phagocytes as antigen presenting cells and as active inducers of thymocyte death. We demonstrate that phagocytosis promotes autoreactive thymocyte death, and that the recognition of PS is critical for this process. Additionally, we show that negative selection is most efficient when the phagocyte also presents the negative selecting peptide. Taken together, our data support a multi-step model for negative selection in which a strong TCR signal initiates apoptosis, followed by phagocytosis to deliver the lethal hit. Moreover, the coupling of these two steps that occurs when phagocytes also serve as APCs leads to more efficient negative selection (Supplementary Fig. 5a).

*In vitro* studies of apoptosis support a model in which a death signal initiates a cell-autonomous program of cellular destruction involving protease and nuclease activation, PS exposure on the outer membrane, membrane blebbing, and the formation of apoptotic bodies^2^. Our current work adds to a growing body of evidence that the early events associated with apoptosis are not necessarily a death sentence. For example, activated T cells can transiently express active caspase-3 and expose PS at the cell surface, but ultimately avoid cell death^31–35^. Moreover, phagocyte-dependent killing is critical for regulating the size of T cell, erythrocyte, and neutrophil populations, as well as removal of transient structures during development^36–44^. Thus, mounting evidence suggests that phagocytosis is a critical final step in cell death *in vivo*.

The mechanism by which phagocytes actually kill their target cells remains unknown. Following phagocytosis, the engulfed cell would be exposed to lysosomal proteases and other degradative enzymes. Furthermore, under certain conditions phagocytes release reactive oxygen species (ROS) into the phagosome, and ROS have been shown to be important for the phagocytosis-induced death of neurons during development in the mouse brain^36^. It is tempting to speculate that the cytotoxic environment that a cell encounters within the phagosome following engulfment is ultimately responsible for its death. Although indirect mechanisms such as retinoic acid and cytokine production by phagocytes^45^ and PS signaling^31^, might also impact thymocyte survival, the fact that we observe similar defects in negative selection upon phagocyte depletion, blocking PS, and mutation of a PS receptor, strongly suggests that phagocytes mediate thymocyte death directly via phagocytosis.

Our observation that negative selection is most efficient when phagocytes also serve as peptide-presenting cells implies that the phagocytic activity of a peptide-presenting cell is a key factor in determining thymocyte fate. Indeed, the inability to phagocytose could help to explain the relative inefficiency of thymic stromal cells to induce negative selection^5–7,29^, in contrast to the phagocytic activity of many hematopoietic cells which are potent inducers of negative selection. It is worth noting that many thymic phagocytes, identified in this study by the expression of the LysM-GFP reporter, also express the DC marker CD11c^13^ (Supplementary Fig. 1). Thus, a subset of thymic hematopoietic cells with characteristics of both DCs and macrophages may be particularly important mediators of negative selection. Although some non-phagocytic subsets, including mTECs, thymocytes, and B cells, can induce negative selection^3,21,46^, this process may be rendered less efficient by the requirement for a second cellular encounter with a phagocyte in order to complete negative selection (Supplementary Fig. 5b).

The initiation of phagocytosis is dependent on the recognition of target cells via a variety of receptors for “eat me” signals, including PS. We found that mutation of a single PS receptor, Tim-4, impaired negative selection to TRA, but had a less obvious impact on negative selection to ubiquitous antigen. On the other hand, globally blocking PS significantly impaired both forms of negative selection. These results could indicate that Tim-4 plays a non-redundant role in phagocytosis in the medulla, the site of TRA presentation, whereas other PS receptors may participate in phagocytosis in the thymic cortex, where negative selection to ubiquitous antigen takes place^5,8,47^. Alternatively, negative selection to TRA might be more sensitive to perturbations in phagocytosis, given that TRA are typically expressed at low levels, and may deliver a relatively weak apoptotic signal. In line with this idea, cells receiving a weak apoptotic signal in *C. elegans* embryos are more dependent upon phagocytosis for their death than cells receiving a strong apoptotic signal^39^. This raises the intriguing possibility that phagocytes might be especially critical during negative selection to relatively low-affinity or rare self-antigens, which pose the greatest risk as targets of autoimmunity^48^.

A requirement for Tim-4 in negative selection involving relatively weak apoptotic stimuli is consistent with the mild autoimmune phenotype reported in Tim-4^−/−^ mice^49^. Although their hyperimmune phenotype was initially attributed to defective phagocytosis in the periphery^44,49^, our data suggest that an increase in the release of autoreactive T cells from the thymus due to defective thymic phagocytes might also contribute. Notably, Tim-4^−/−^ mice do not develop overt autoimmunity, likely due to the presence of other tolerance mechanisms, such as regulatory T cells and other peripheral tolerance mechanisms, which could serve to keep self-reactive T cells in check.

Our current data contribute to emerging evidence that the context in which an autoreactive thymocyte encounters peptides, shaped largely by the characteristics of the peptide-presenting cell, has profound impacts on its fate. We recently reported that thymic dendritic cells that provide both high-affinity TCR ligands and a local source of IL-2 can efficiently support the development of regulatory T cells^30,47^. Thus, the decision of an autoreactive thymocyte to die or differentiate may ultimately depend on whether it engages a peptide-presenting cell that promotes its death or supports its further development.

## Materials and methods

### Mice

All mice were bred and maintained under pathogen-free conditions in an American Association of Laboratory Animal Care-approved facility at the University of California, Berkeley. The University of California, Berkeley Animal Use and Care Committee approved all procedures. C57BL/6, C57BL/6-Tg(Ins2-TFRC/OVA)296Wehi/WehiJ (RIPmOVA), and C567BL/6-Tg(Csf1r-EGFP-NGFR/FKBP1A/TNFRSF6)2Bck/J (MAFIA) mice were from Jackson Labs. OT-I Rag2^−/−^ mice were from Taconic Farms. LysMGFP, F5 Rag1^−/−^, and Tim-4^−/−^ mice have been previously described^19,50,51^. LysMGFP RIPmOVA and Tim-4^−/−^ RIPmOVA mice were generated by crossing LysMGFP or Tim-4^−/−^ mice to RIPmOVA mice. Mice were used from four to eight weeks of age.

### Thymocyte Isolation and Labeling

Thymuses were collected from OT-I Rag2^−/−^, F5 Rag1^−/−^, or B6 mice and dissociated through a 70μm cell strainer to yield a cell suspension. Thymocytes were then labeled with 1μM Cell Proliferation Dye eFluor450 or 0.5μM Cell Proliferation Dye eFluor670 (Thermo Fisher Scientific) at 10^7^ cells/ml at 37°C for 15 minutes in PBS, then washed and resuspended in complete RPMI (containing 10% FBS, penicillin streptomycin, and 2-mercaptoethanol, cRPMI) for overlay onto thymic slices. Thymocytes do not proliferate at the timepoints collected, allowing overlaid thymocytes to be distinguished from slice resident thymocytes by Cell Proliferation Dyes (Fig. 1a). In imaging experiments, OT-I thymocytes were depleted of mature CD8 single positives using the EasySep Biotin Positive Selection Kit (Stemcell Technologies) with anti-human/mouse β7 integrin antibody (FIB504, Biolegend) according to the manufacturer’s instructions. Thymocytes were then labeled with 3μM SNARF (Thermo Fisher Scientific) at 10^7^ cells/ml at 37°C for 15 minutes in PBS, then washed and labeled with 5μM Hoechst 33342 (Thermo Fisher Scientific) at 10^7^ cells/ml at 37°C for 15 minutes.

### Thymic Slices

Preparation of thymic slices has been previously described^52,53^. Thymic lobes were cleaned of connective tissue, embedded in 4% agarose with a low melting point (GTG-NuSieve Agarose, Lonza), and sectioned into slices of 200-400μm using a vibratome (1000 Plus sectioning system, Leica). Slices were overlaid onto 0.4μm transwell inserts set in 6 well tissue culture plates with 1ml cRPMI under the insert. 0.5-2 ×10^6^ thymocytes in 10μl cRPMI were overlaid onto each slice and allowed to migrate into the slice for 2 hours, then excess thymocytes were removed by gentle washing with PBS. Thymocytes actively migrate into the slice and localize as expected based on their maturation status^24,26,54^. For peptide-induced negative selection, 10μl of 1μM SIINFEKL (AnaSpec) in PBS was overlaid onto each slice for 30 minutes, then removed by pipetting. To quantify negative selection, we used a fluorescent live/dead stain (Ghost Dye Violet 510 or Fixable Viability Dye eFluor780) to identify live cells (as shown in Fig. 1a). We then calculated the ratio of total live OT-I thymocytes to total live reference (either wild type thymocytes or thymocytes expressing an irrelevant TCR: F5) recovered from the thymic slice. In general, the ratio of OT-I to reference populations was close to 1.0 in the absence of OVA, however, there was some variability due to differential ability of the two populations to enter or survive in the tissue. We therefore further normalized the ratios of OT-I:reference thymocytes in each experiment so that the average of the corresponding “no OVA” samples was always 1.0. For depletion of phagocytes, 1μM AP20187 (Clontech) was added to the media under the transwell and 10μl of 10μM AP20187 in PBS was added on top of each slice overnight (16-18 hours). The drug was washed out from the top of the slice with PBS prior to overlaying thymocytes. For Annexin V treatment, thymocytes were resuspended in Annexin V binding buffer (Thermo Fisher Scientific) with purified Annexin V (BioLegend) at 200μg/ml prior to overlaying on the slice. Following peptide treatment, 10μl of purified Annexin V at 200μg/ml in Annexin V binding buffer was overlaid onto each slice.

### Bone marrow-derived dendritic cell cultures

Bone marrow was flushed from the femurs and tibias of mice into sterile PBS, and treated with ammonium chloride–potassium bicarbonate buffer for lysis of red blood cells. Cells were resuspended at 10^6^/ml in cRPMI with 20ng/ml granulocyte-macrophage colony-stimulating factor (GM-CSF, Peprotech) and plated for culture. Cells were cultured for 7 days, with replacement with fresh media containing GM-CSF on day 6. On day 7, semi-adherent cells were collected and loaded with 1μM SIINFEKL at 10^7^/ml in cRPMI at 37°C for 30 minutes. Some BMDCs were incubated without peptide, as indicated. Cells were then washed and 10^5^ BMDCs were overlaid per slice, following washout of excess thymocytes.

### Flow cytometry

Thymic slices, whole thymuses, and spleens were dissociated into FACS buffer (0.5% BSA in PBS) and filtered before staining. Splenocytes and blood samples were treated with ammonium chloride-potassium bicarbonate buffer for 5 or 10 minutes, respectively, at room temperature prior to staining to lyse red blood cells. Cells were stained for 10 minutes on ice in 24G2 supernatant containing the following antibodies: CD4 (GK1.5), CD8α (53-6.7), CD69 (H1.2F3). Cells were then washed in PBS and stained in Ghost Dye Violet 510 (Tonbo Biosciences) or Fixable Viability Dye eFluor780 (Thermo Fisher Scientific) for 10 minutes on ice. For staining of thymic phagocyte populations, whole thymuses were minced and incubated in cRPMI containing 1mg/ml collagenase Type IA (Sigma) and 400μg/ml DNase I (Roche) at 37°C for 1 hour. After vigorous pipetting, samples were filtered, then stained in 24G2 supernatant containing the following antibodies: CD11b (M1/70), CD11c (N418), F4/80 (BM8), Tim-4 (RMT4-54), MHC I H-2Kd/H2-Dd (34-1-2S), MHC II I-A/I-E (M5/114.15.2), CD80 (16-10A1), CD86 (GL1), ICAM (YN1/1.7.4). Cells were then washed in PBS and stained in Fixable Viability Dye eFluor780 (Thermo Fisher Scientific) for 10 minutes on ice. All antibodies were from Thermo Fisher Scientific, Biolegend, or Tonbo Biosciences. Flow cytometry was performed with a LSRII or Fortessa X20 (BD Biosciences) and FlowJo software (TreeStar) was used for data analysis. Gating strategies are shown for thymocytes (Fig. 1a) and thymic phagocyte populations (Supplementary Fig. 1).

### Two-photon microscopy

Two-photon imaging of thymic slices has been described previously^10,29,52^. Briefly, thymic slices were glued to coverslips and fixed to a dish being perfused at a rate of 1ml/minute with oxygenated, phenol red-free DMEM media warmed to 37°C. Imaging was performed with a Zeiss 7 MP two-photon microscope with a Coherent Chameleon laser tuned to 920nm. Signals were separated using 495nm and 560nm dichroic mirrors. Imaging volumes were scanned every 30 seconds for 30 minutes, and images were processed with Imaris 7.3 software (Bitplane).

### Statistics

Statistical analysis was carried out using Prism software (GraphPad). The D’Agostino and Pearson omnibus K2 normality test, Shapiro-Wilk normality test, or Kolmogorov-Smirnov test was applied depending on sample size, and parametric or non-parametric statistical analyses were carried out as appropriate (specific tests used are indicated in figure legends). P values of <0.05 were considered significant.

## Acknowledgements

We would like to thank members of the Robey lab for technical advice and helpful discussion, H. Melichar, B. Weist, and B.J. Fowlkes for critical reading of the manuscript, S. W. Chan and O. Guevarra for technical assistance, P. Herzmark for help with two-photon imaging, and Wenjun Ouyang (Genentech) for providing Tim4^−/−^ mice. Supported by the National Institutes of Health (R01AI064227 to E.A.R.) and a University of California Cancer Research Coordinating Committee Fellowship to N.S.K..

