## Supplemental Figures for "A role for phagocytosis in inducing cell death during thymocyte negative selection"

### Supplementary Figures

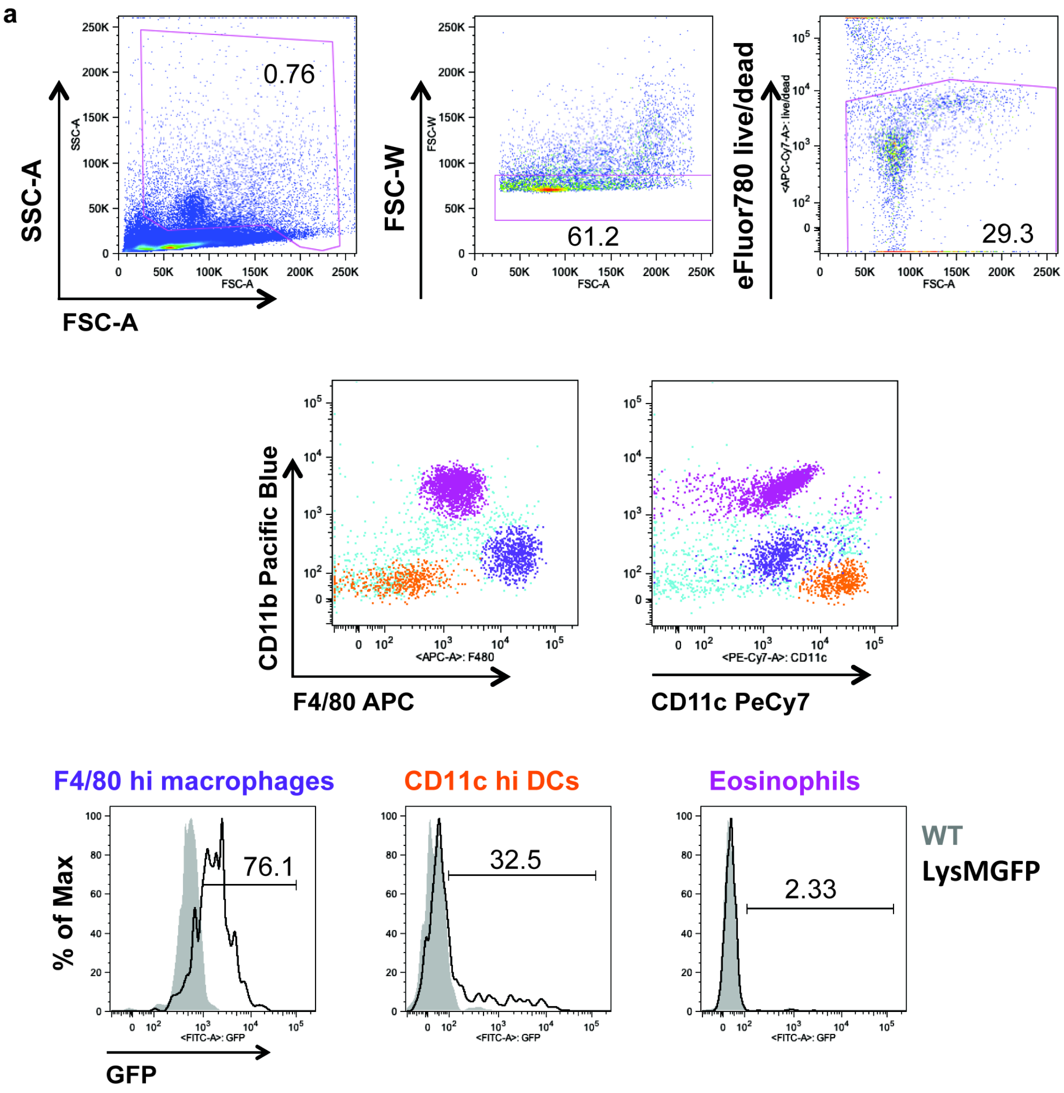


**Supplementary Figure 1. GFP-expressing cells in the LysMGFP thymus include F4/80$^{hi}$ macrophages as well as a subset of CD11c$^{hi}$ DCs**

(a) Major populations of thymic myeloid cells (gated on live singlets with high forward and side scatter) include F4/80$^{hi}$ macrophages (purple), CD11c$^{hi}$ DCs (orange), and F4/80-intermediate CD11c-intermediate eosinophils (magenta). Gating strategy shown was used for all experiments where thymic myeloid cells were analyzed. (b) Expression of GFP in thymic myeloid cells in LysMGFP mice (black), with WT controls shown in gray. Data are representative of n= 2 mice per condition.

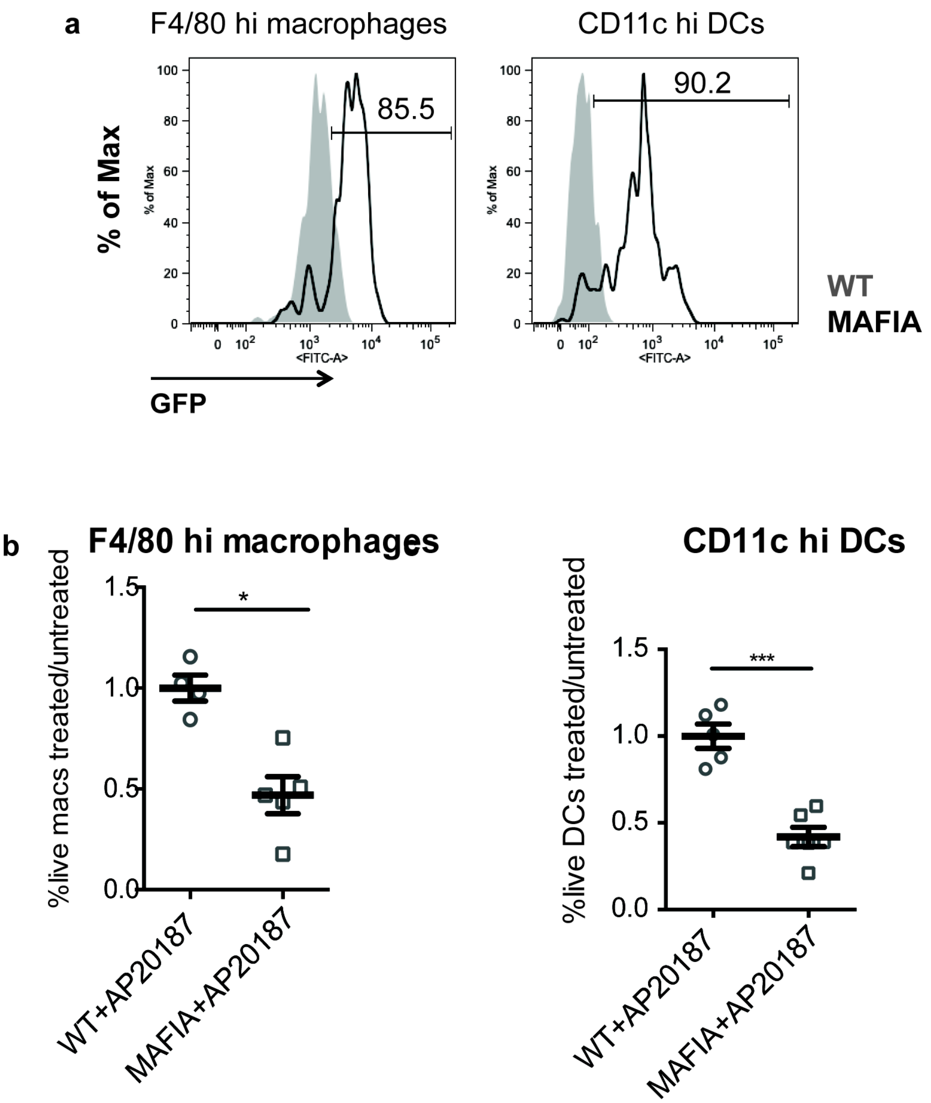


**Supplementary Figure 2. Depletion of phagocytes in MAFIA thymic slices**
(a) Expression of the MAFIA transgene in F4/80$^{hi}$ macrophages (left) and CD11c$^{hi}$ DCs (right). (b,c) WT or MAFIA thymic slices were treated with AP20187 or left untreated for 16-18 hours. Slices were then dissociated and proportions of live F4/80$^{hi}$ macrophages (b) or CD11c$^{hi}$ DCs (c) remaining in the slice were determined by flow cytometry. Fold depletion of macrophages (b) or DCs (c) upon treatment displayed as the proportion of live phagocytes present in treated slices relative to untreated slices. Data are representative of 3 independent experiments, with mean and SEM of n=4 (WT F4/80$^{hi}$ macrophages), 5 (MAFIA F4/80$^{hi}$ macrophages, WT CD11c$^{hi}$ DCs), or 6 (MAFIA CD11c$^{hi}$ DCs) total slices shown. ns not significant ($p>0.05$), *$p<0.05$, ***$p<0.001$ (two-tailed unpaired Student's *t*-test with 95% confidence interval).

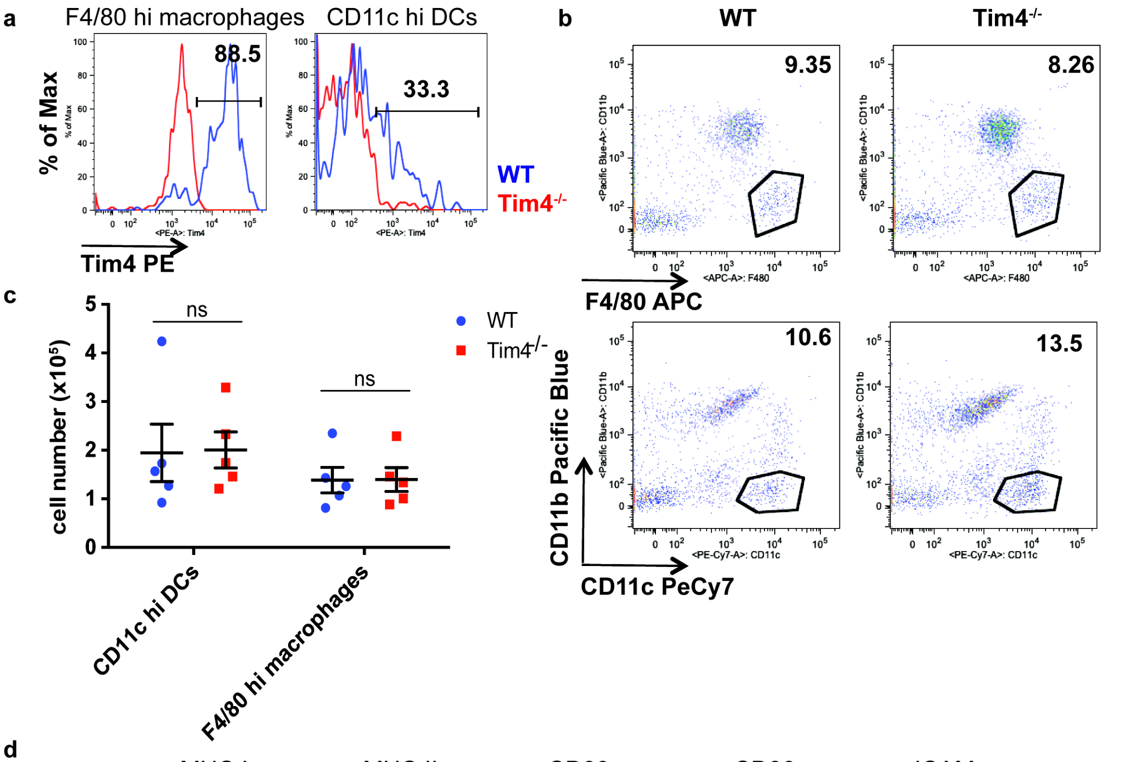


**Supplementary Figure 3. Normal number and cell-surface phenotype of phagocytes in the thymus of Tim-4$^{-/-}$ mice**

(a) Expression of Tim-4 by WT F4/80$^{hi}$ macrophages (left) and CD11c$^{hi}$ DCs (right) shown in blue, with Tim-4$^{-/-}$ shown in red for comparison. (b,c) Proportions (b) or total number per thymus (c) of macrophages and DCs in age and sex-matched WT and Tim-4$^{-/-}$ mice. Data are representative of (a,b) or compiled from (c) n=5 mice per genotype, where each dot represents an individual mouse. ns not significant ($p>0.05$) (two-tailed Mann-Whitney test, macrophages, or Student's *t*-test, DCs, test with 95% confidence interval) (d) Expression of markers associated with antigen presentation and costimulation by WT (blue) and Tim-4$^{-/-}$ thymic F4/80 hi macrophages and CD11c hi DCs. Data are representative of three independent experiments.

a

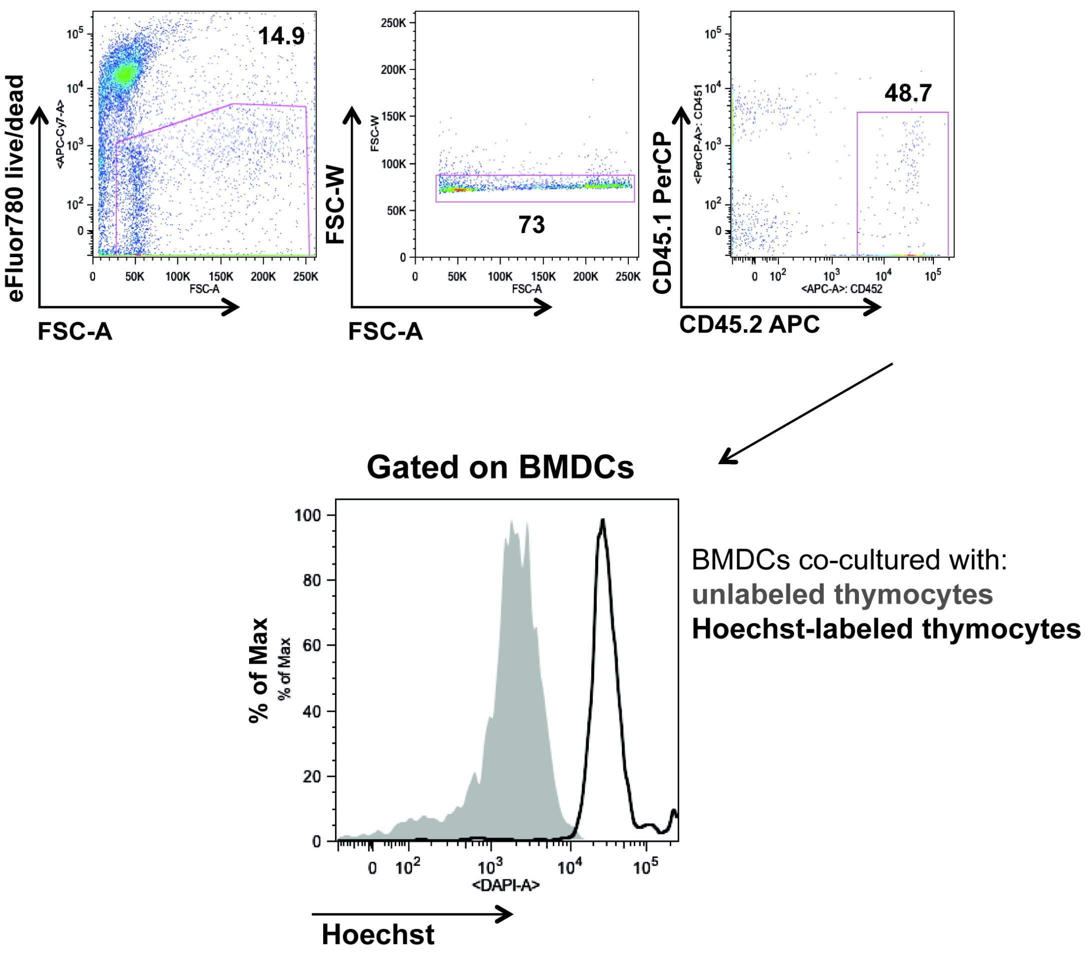


**Supplementary Figure 4. Bone marrow-derived dendritic cells are phagocytic**
Hoechst-labeled or unlabeled CD45.1 OT-I thymocytes were cultured in vitro for 16 hours with CD45.2 BMDCs. Accumulation of Hoecsht dye in BMDCs (gated on live, CD45.2+, singlets) cultured with unlabeled (gray) or Hoecsht-labeled (black) thymocytes.

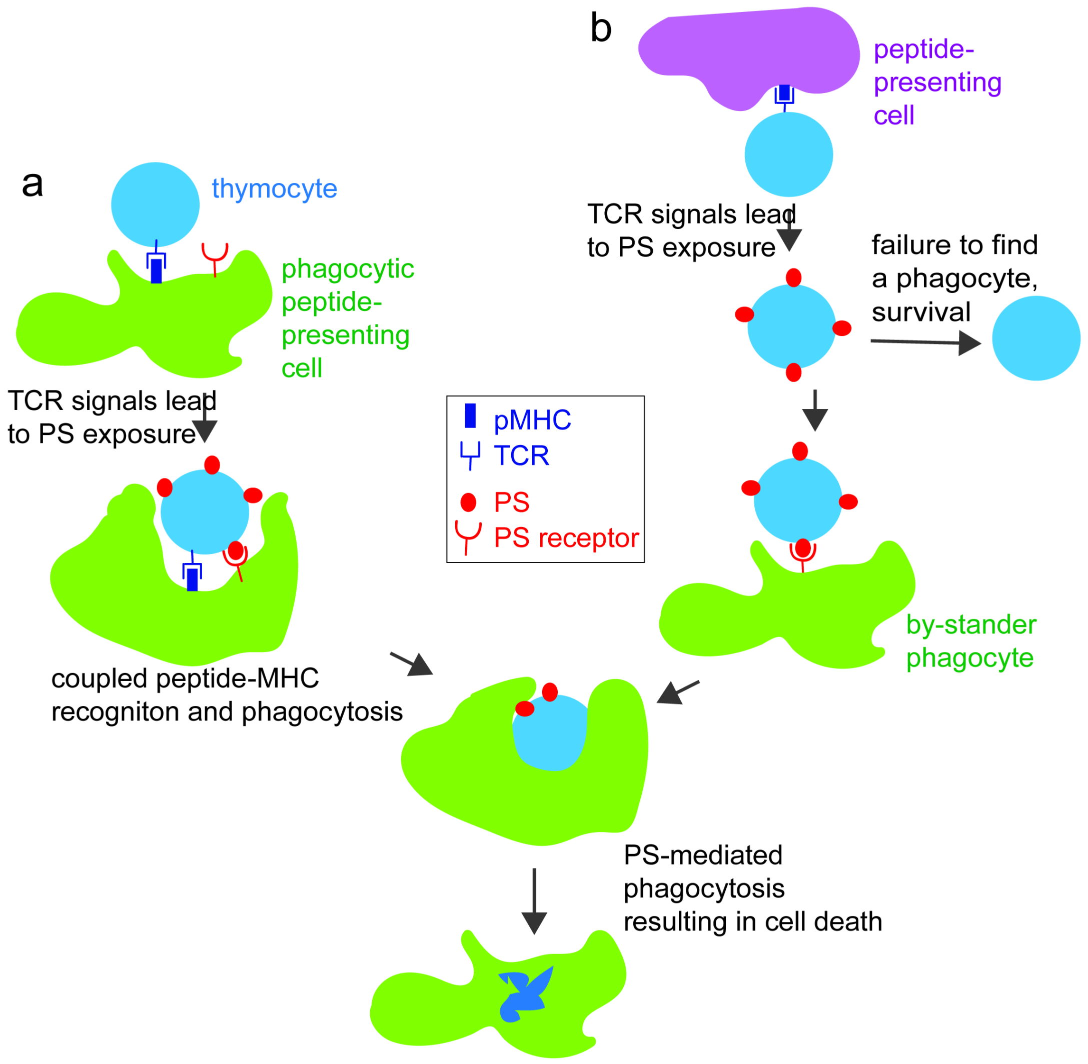


**Supplementary Figure 5. Model for the role of phagocytes in inducing cell death during negative selection**

A two-step model for negative selection. In the first step, a thymocyte encounters a high affinity self-peptide-MHC ligand (pMHC, dark blue rectangles) on a peptide presenting cell, leading to the exposure of phosphatidyl serine (PS, red ovals) on the cell surface. In a second step, recognition of the autoreactive thymocyte via PS-receptors on a phagocyte leads to uptake and death of the thymocyte. In (a) the same cell both presents the self-peptide and phagocytoses the autoreactive thymocyte, leading to efficient negative selection. In (b), the thymocyte encounters peptide on a non-phagocytic cell, and requires a subsequent interaction with a phagocyte in order to mediate its death. Thymocytes that fail to subsequently interact with a phagocyte may recover and survive, leading to less efficient negative selection.

**Supplementary movies**

Examples of thymocyte death occurring concurrently with phagocytosis. OT-I thymocytes were depleted of mature CD8 SP and double labeled with Hoechst and SNARF before overlaying on LysMGFP RIPmOVA thymic slices. Slices were imaged by two-photon scanning laser microscopy at 9.5 hours after thymocyte addition to the slice.
